# Detection and interpretation of shared genetic influences on 40 human traits

**DOI:** 10.1101/019885

**Authors:** Joseph K. Pickrell, Tomaz Berisa, Laure Segurel, Joyce Y. Tung, David Hinds

## Abstract

We performed a genome-wide scan for genetic variants that influence multiple human phenotypes by comparing large genome-wide association studies (GWAS) of 40 traits or diseases, including anthropometric traits (e.g. nose size and male pattern baldness), immune traits (e.g. susceptibility to childhood ear infections and Crohn’s disease), metabolic phenotypes (e.g. type 2 diabetes and lipid levels), and psychiatric diseases (e.g. schizophrenia and Parkinson’s disease). First, we identified 307 loci (at a false discovery rate of 10%) that influence multiple traits (excluding “trivial” phenotype pairs like type 2 diabetes and fasting glucose). Several loci influence a large number of phenotypes; for example, variants near the blood group gene ABO influence eleven of these traits, including risk of childhood ear infections (rs635634: log-odds ratio = 0.06, P = 1.4 *×* 10^−8^) and allergies (log-odds ratio = 0.05, *P* = 2.5 *×* 10^−8^), among others. Similarly, a nonsynonymous variant in the zinc transporter SLC39A8 influences seven of these traits, including risk of schizophrenia (rs13107325: log-odds ratio = 0.15, *P* = 2 *×* 10^−12^) and Parkinson’s disease (log-odds ratio = -0.15, *P* = 1.6 *×* 10^−7^), among others. Second, we used these loci to identify traits that share multiple genetic causes in common. For example, genetic variants that delay age of menarche in women also, on average, delay age of voice drop in men, decrease body mass index (BMI), increase adult height, and decrease risk of male pattern baldness. Finally, we identified four pairs of traits that show evidence of a causal relationship. For example, we show evidence that increased BMI causally increases triglyceride levels, and that increased liability to hypothyroidism causally decreases adult height.

## 1 Introduction

The observation that a genetic variant affects multiple phenotypes (a phenomenon often called “pleiotropy” [Paaby and Rockman, 2013; Solovieff et al., 2013; Stearns, 2010], though we will not use this term) is informative in a number of applications. One such application is to learn about the molecular function of a gene. For example, men with the genetic disease cystic fibrosis (primarily known as a lung disease) are often infertile due to congenital absence of the vas deferens; this is evidence of a shared role for the CFTR protein in lung function and the development of reproductive organs [Chillón et al., 1995]. Another application is to learn about the causal relationships between traits. For example, individuals with congenital hypercholesterolemia also have elevated risk of heart disease [Müller, 1938]; this is now interpreted as evidence that changes in lipid levels causally influence heart disease risk [Steinberg, 2002].

In these two applications, the same observation–that a genetic variant influences two traits–is interpreted in fundamentally different ways depending on known aspects of biology. In the first case, a genetic variant influences the two phenotypes through independent physiological mechanisms (graphically: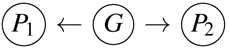, if *G* represents the genotype, *P*_1_ the first phenotype, *P*_2_ the second phenotype, and the arrows represent causal relationships [Pearl, 2000]), while in the second case, 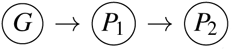. In some situations, knowing which interpretation of the observation to prefer is simple: for example, it seems difficult to imagine how the reproductive and lung phenotypes of a CFTR mutation could be related in a causal chain. In other situations, interpretation is considerably more challenging. For example, the causal connections between various lipid phenotypes and heart disease have been debated for decades (e.g. Steinberg [1989]).

As the number of reliable associations between genetic variants and various phenotypes has grown over the last decade [Visscher et al., 2012], these issues have received increasing attention. A number of studies have identified genetic variants that influence multiple traits [Andreassen et al., 2013a,b; Cotsapas et al., 2011; Elliott et al., 2013; Estrada et al., 2012; Li et al., 2014; Moltke et al., 2014; Pendergrass et al., 2013; Sivakumaran et al., 2011; Stefansson et al., 2014; Styrkarsdottir et al., 2013]; in general, these associations are interpreted as most plausibly due to independent effects of a genetic variant on different aspects of physiology. For example, a genetic variant in LGR4 is associated with bone mineral density (BMD), age at menarche, and risk of gallbladder cancer [Styrkarsdottir et al., 2013], presumably due to effects mediated through different tissues.

There has also been increasing interest in the alternative, causal framework for interpreting genetic variants that influence multiple phenotypes, which has been formalized under the name “Mendelian randomization” [Davey Smith and Ebrahim, 2004; Davey Smith and Hemani, 2014; Katan, 1986]. Mendelian randomization has been used to provide evidence for (or against) a causal role for various clinical variables in disease etiology [De Silva et al., 2011; Granell et al., 2014; Holmes et al., 2014; Lim et al., 2014; Panoutsopoulou et al., 2013; Pichler et al., 2013; Voight et al., 2012]. For example, genetic variants associated with body mass index (BMI) are also associated with type 2 diabetes [Holmes et al., 2014]; this is consistent with a causal role for weight gain in the etiology of diabetes.

To date, most studies of multiple traits have been performed in a targeted fashion–for example, there have been scans for variants that influence multiple autoimmune diseases [Cotsapas et al., 2011] or multiple psychiatric phenotypes [Cross-Disorder Group of the Psychiatric Genomics Consortium, 2013]. We aimed to systematically search for genetic variants that influence pairs of traits, and then to interpret these associations in the light of the causal and non-causal models described above. In this paper, we describe the results of such a search using large genome-wide association studies of 40 traits.

## 2 Results

We assembled summary statistics from 41 genome-wide association studies of 40 traits or diseases performed in individuals of European descent (Table 1; two of these GWAS are for age at menarche). These studies span a wide range of phenotypes, from anthropometric traits (e.g. height, BMI, nose size) to neurological disease (e.g. Alzheimer’s disease, Parkinson’s disease) to susceptibility to infection (e.g. childhood ear infections, tonsillectomy). For studies that were not done using imputation to all variants in phase 1 of the 1000 Genomes Project [Abecasis et al., 2010], we performed imputation at the level of summary statistics using ImpG v1.0 [Pasaniuc et al., 2014]. We estimated the approximate number of independent associated variants (at a false discovery rate of 10%) in each study using fgwas v.0.3.6 [Pickrell, 2014]. The number of associations ranged from around five (for age at voice drop in men) to over 500 (for height).

**Table 1.** Phenotypes used in this study. For each study, we show the name of the phenotype, the abbreviation that will be used throughout this paper, the data source, the number of independent autosomal loci identified at a false discovery rate of 10%, and the number of participants in the study. For studies where the data source is 23andMe, a complete description of the GWAS is presented in the Supplementary Material.

| Phenotype | Abbreviation | Data source | Approx. # of loci | Approx. # of participants, in thousands (cases/controls, if applicable) |
| --- | --- | --- | --- | --- |
| Alzheimer's disease | AD | [Lambert et al., 2013] | 11 | 17 / 37 |
| Age at menarche | AAM | [Perry et al., 2014] | 70 | 133 |
| Age at menarche (23andMe) | AAM (23) | 23andMe | 55 | 77 |
| Height | HEIGHT | [Wood et al., 2014] | 584 | 253 |
| Schizophrenia | SCZ | [Psychiatric Genomics Consortium, 2014] | 222 | 34 / 46 |
| Rheumatoid arthritis | RA | [Okada et al., 2014] | 74 | 14 / 44 |
| Coronary artery disease | CAD | [Schunkert et al., 2011] | 11 | 22 / 65 |
| Type 2 diabetes | T2D | [Morris et al., 2012] | 11 | 12 / 57 |
| Crohn's disease | CD | [Jostins et al., 2012] | 61 | 6 / 15 |
| Fasting glucose | FG | [Manning et al., 2012] | 15 | 58 |
| Hemoglobin | HB | [van der Harst et al., 2012] | 16 | 51 |
| Mean cell hemoglobin concentration | MCHC | [van der Harst et al., 2012] | 15 | 46 |
| Mean red cell volume | MCV | [van der Harst et al., 2012] | 42 | 48 |
| Packed red cell volume | PCV | [van der Harst et al., 2012] | 13 | 44 |
| Red blood cell count | RBC | [van der Harst et al., 2012] | 25 | 45 |
| Body mass index | BMI | [Locke et al., 2015] | 30 | 240 |
| Waist-hip ratio | WHR | [Shungin et al., 2015] | 13 | 143 |
| Low-density lipoproteins | LDL | [Teslovich et al., 2010] | 41 | 85 |
| High-density lipoproteins | HDL | [Teslovich et al., 2010] | 46 | 89 |
| Triglycerides | TG | [Teslovich et al., 2010] | 31 | 86 |
| Total cholesterol | TC | [Teslovich et al., 2010] | 53 | 89 |
| Bone mineral density (femoral neck) | FNBMD | [Estrada et al., 2012] | 19 | 33 |
| Bone mineral density (lumbar spine) | LSBMD | [Estrada et al., 2012] | 21 | 32 |
| Platelet count | PLT | [Gieger et al., 2011] | 50 | 44 |
| Mean platelet volume | MPV | [Gieger et al., 2011] | 29 | 17 |
| Any allergies | ALL | 23andMe | 43 | 67 / 114 |
| Asthma | ATH | 23andMe | 35 | 28 / 129 |
| Age at voice drop | AVD | 23andMe | 5 | 56 |
| Beighton hypermobility | BHM | 23andMe | 18 | 64 |
| Childhood ear infections | CEI | 23andMe | 15 | 47 / 75 |
| Breast size | CUP | 23andMe | 14 | 34 |
| Chin dimples | DIMP | 23andMe | 57 | 58 / 13 |
| Hypothyroidism | HTHY | 23andMe | 30 | 18 / 117 |
| Migraine | MIGR | 23andMe | 37 | 53 / 231 |
| Male pattern baldness | MPB | 23andMe | 49 | 9 / 8 |
| Nose size | NOSE | 23andMe | 13 | 67 |
| Nearsightedness | NST | 23andMe | 183 | 106 / 86 |
| Parkinson's disease | PD | 23andMe | 43 | 10 / 325 |
| Photoc sneeze reflex | PS | 23andMe | 66 | 32 / 67 |
| Tonsillectomy | TS | 23andMe | 48 | 60 / 113 |
| Unibrow | UB | 23andMe | 61 | 69 |

### 2.1 A model for identification of genetic variants that influence pairs of traits

We first aimed to identify genetic variants that influence pairs of traits. To do this, we developed a statistical model (extending that used by Giambartolomei et al. [2014]) to estimate the probability that a given genomic region either 1) contains a genetic variant that influences the first trait, 2) contains a genetic variant that influences the second trait, 3) contains a genetic variant that influences both traits, or 4) contains both a genetic variant that influences the first trait and a separate genetic variant that influences the second trait (Figure 1). The input to the model is the set of summary statistics (effect size estimates and standard errors) for each SNP in the genome on each of the two phenotypes, and (if the two GWAS were performed on overlapping sets of individuals) the expected correlation in the summary statistics due to correlation between the phenotypes. We can then fit the following log-likelihood function:

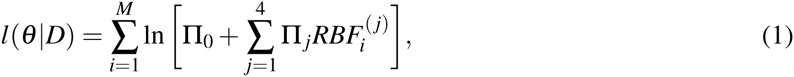

where *D* is the data, *M* is the number of approximately independent blocks in the genome, Π_0_ is the prior probability that a region contains no genetic variants than influence either trait, Π_1_, Π_2_, Π_3_, and Π_4_ represent the prior probabilities of the four models described above, *θ* is the set of all five prior parameters, and 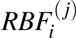 is the regional Bayes factor measuring the support for model *j* in genomic region *i* (see Methods for details). In the presence of missing data, we consider only the subset of SNPs with data in both studies; if the causal SNP is not present this acts to reduce power to detect a shared effect [Giambartolomei et al., 2014]. In fitting this model, we estimate the prior parameters and the posterior probability of each model for each region of the genome (for numerical stability, in practice we penalize the estimates of the prior parameters, and so obtain maximum *a posteriori* estimates). We were mainly interested in the estimated prior probability that each genomic region contains a variant that influences both trait 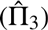 and the corresponding posterior probabilities for each genomic region.

**Figure 1.**
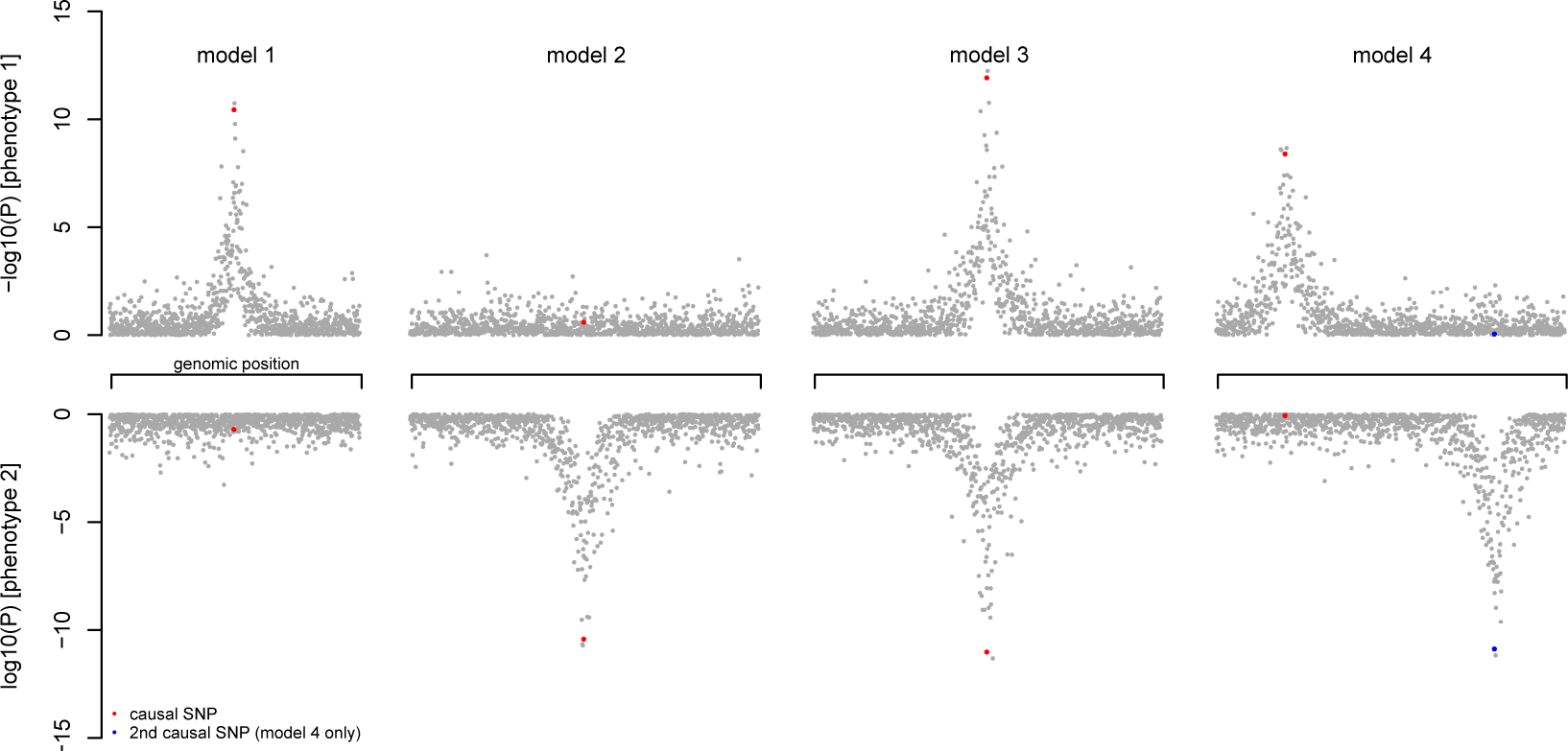
Schematic of the different models considered for a given genomic region and two GWAS. We divide the genome into approximately independent blocks (see Methods), and estimate the proportion of blocks that fit into the shown patterns. The null model with no associations is not shown. Each point represents a single genetic variant.

Several caveats of this method are worth mentioning. First, note that the parameter Π_3_ is best thought of as the proportion of genomic regions that *detectably* influence both traits–if one study is small and underpowered, this estimate will necessary be zero. This contrasts with methods that aim to provide unbiased estimates of the “genetic correlation” between traits that do not depend on sample size [Bulik-Sullivan et al., 2015; Loh et al., 2015; Yang et al., 2011]. Second, in general it is not possible to distinguish a single causal variant that influences both traits (Model 3 in Figure 1) from two separate causal variants (Model 4 in Figure in the presence of strong linkage disequilibrium between the causal variants. For any individual genomic region discussed below, the possibility of two highly correlated causal variants must be considered as an alternative possibility in the absence of functional follow-up. Finally, we evaluated the method in simulations (Supplementary Figures 1-4), and found that the model gives a small overestimate of proportion of shared effects (Supplementary Figure 3). This is because the amount of evidence against the null model of no associations is greater when a variant influences both phenotypes compared to when it only influence a single phenotype (Supplementary Figure 4).

### 2.2 Identification of variants that influence pairs of traits across 41 GWAS

We applied the method to all pairs of the 41 GWAS listed in Table 1. For each pair of studies, we first estimated the expected correlation in the effect sizes from the summary statistics, and included this correction for overlapping individuals in the model. Note that this is conservative: in pairs of GWAS where we are sure there are no overlapping individuals (for example, age at menarche and age at voice drop) we see that the correlation in the summary statistics is non-zero, indicating that we are correcting out some truly shared genetic effects on the two traits (Supplementary Figure 6).

To gain an exploratory sense of the relationships between the phenotypes, we examined the patterns of overlap in associations among all 41 studies. Specifically, the model can be used to estimate, for each pair of traits [*i, j*], the proportion of detected variants that influence trait *i* that also detectably influence trait *j*. These estimates are shown in Figure 2, with phenotypes clustered according to their patterns of overlap. We see several clusters of related traits. For example, of the variants that detectably influence age at menarche (in the Perry et al. [2014] study), the maximum *a posteriori* estimate is that 36% detectably influence height, 30% detectably influence age at voice drop, 28% influence BMI, 10% influence breast size, and 10% influence male pattern baldness. We interpret this as a set of phenotypes that share hormonal regulation. Additionally, there is a large cluster of phenotypes including coronary artery disease, type 2 diabetes, red blood cell traits, and lipid traits, which we interpret as a set of metabolic traits. Further, immune-related disease (allergies, asthma, hypothyroidism, Crohn’s disease and rheumatoid arthritis) all cluster together, and also cluster with infectious disease traits (childhood ear infections and tonsillectomy). This biologically-revelant clustering validates the principle that GWAS variants can identify shared mechanisms underlying pairs of traits in a systematic way. As a control, we performed the same clustering of phenotypes by the estimated proportion of genomic regions where two causal sites fall nearby (Model 4 in Figure 1). In this case, there was no biologically-meaningful clustering (Supplementary Figure 7).

**Figure 2.**
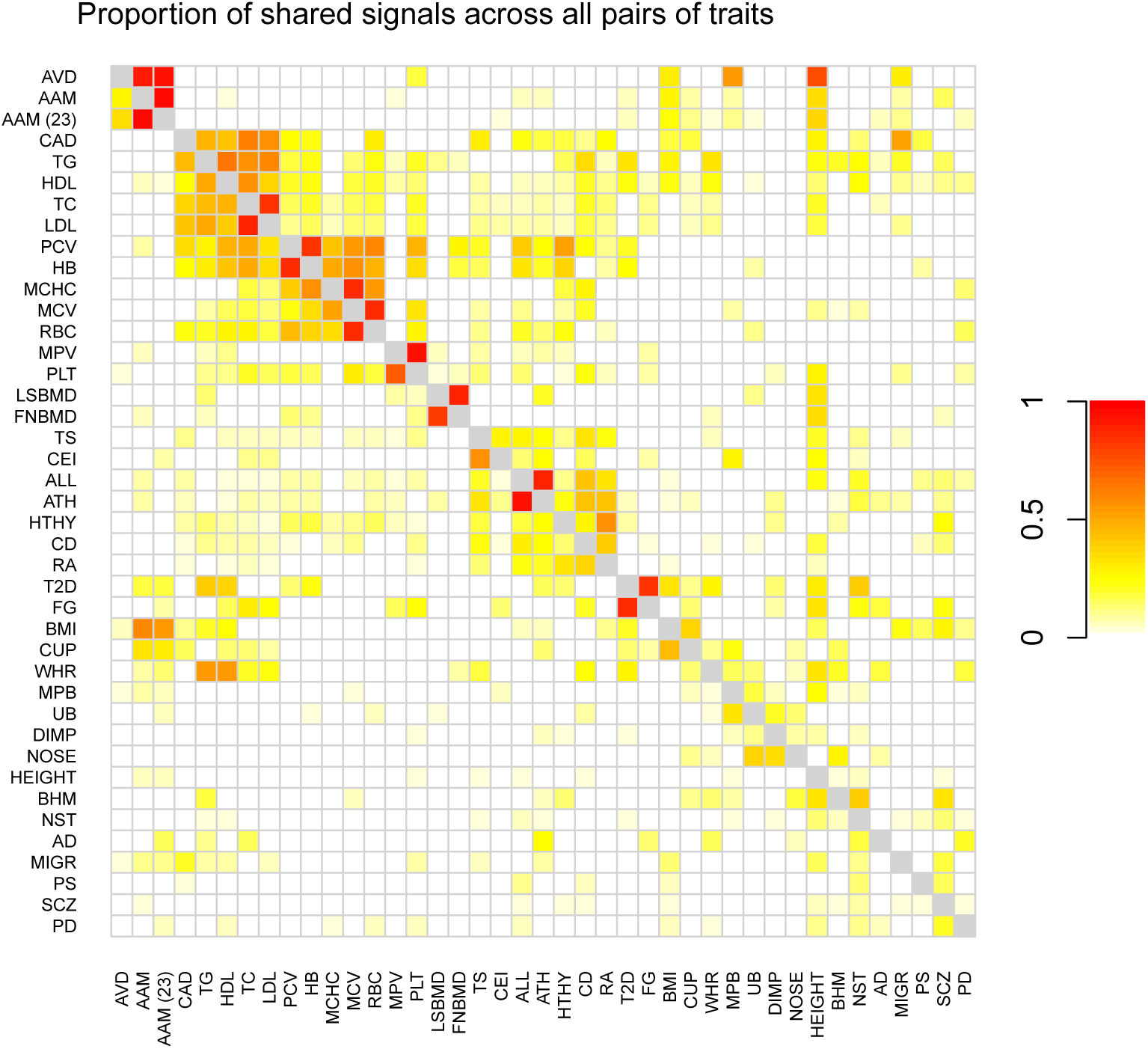
Heatmap showing patterns of overlap between traits. Each square [*i, j*] shows the maximum *a posteriori* estimate of the proportion of genetic variants that influence trait *i* that also influence trait *j*, where *i* indexes rows and *j* indexes columns. Note that this is not symmetric. Darker colors represent larger proportions. Colors are shown for all pairs of traits that have at least one region in the set of 307 identified loci; all other pairs are set to white. Phenotypes were clustered by hierarchical clustering in R [R Core Team, 2013]

### 2.3 Individual loci that influence many traits

We next examined the individual loci identified by these pairwise GWAS. We identified 307 genomic regions where we infer the presence of a variant that influences a pair of traits, at a threshold of a posterior probability greater than 0.9 of model 3 (Supplementary Table 1). This number excludes “trivial” findings where a genetic variant influences two similar traits (two lipid traits, two red blood cell traits, two platelet traits, both measures of bone mineral density, or type 2 diabetes and fasting glucose) and the MHC region.

Some genomic regions contain variants that influence a large number of the traits we considered. We ranked each genomic region according to how many phenotypes share genetic associations in the region (that is, if the pairwise scan for both height and CAD, and the pairwise scan for CAD and LDL, both indicated the same region, we counted this as three phenotypes sharing an association in the region) The top region in this ranking identified a non-synonymous polymorphism in SH2B3 (rs3184504) that is associated with a number of autoimmune diseases, lipid traits, heart disease, and red blood cell traits (Supplementary Figure 8; Supplementary Table 2). This variant has been identified in many GWAS, particularly for autoimmune disease [Richard-Miceli and Criswell, 2012].

The next region in this ranking contains the gene coding for the ABO blood groups in humans, and has a variant associated with 11 traits in these data (and many other additional traits not in these data, see also [Franchini and Lippi, 2015; Schunkert et al., 2011; Wessel et al., 2015]). In Figure 3A, we show the association statistics in this region for coronary artery disease and probability of having a tonsillectomy. At the lead SNP, the non-reference allele is associated with increased risk of CAD (Z = 5.7; P = 1.1 *×* 10^−8^) and increased risk of having a tonsillectomy (Z = 6.0; P = 1.5 *×* 10^−9^). This variant is also strongly associated with other immune, red blood cell, and lipid traits in these data (Figure 3B). A tag for a microsatellite that influences the expression of ABO [Kominato et al., 1997] is correlated to the lead SNP rs635634, as is a tag for the O blood group (Figure 3A). However, the lead SNP is an eQTL for both ABO and the nearby gene SLC2A6 in whole blood [Wessel et al., 2015], so this allele may in fact have downstream effects via effects on the expression of two genes.

**Figure 3.**
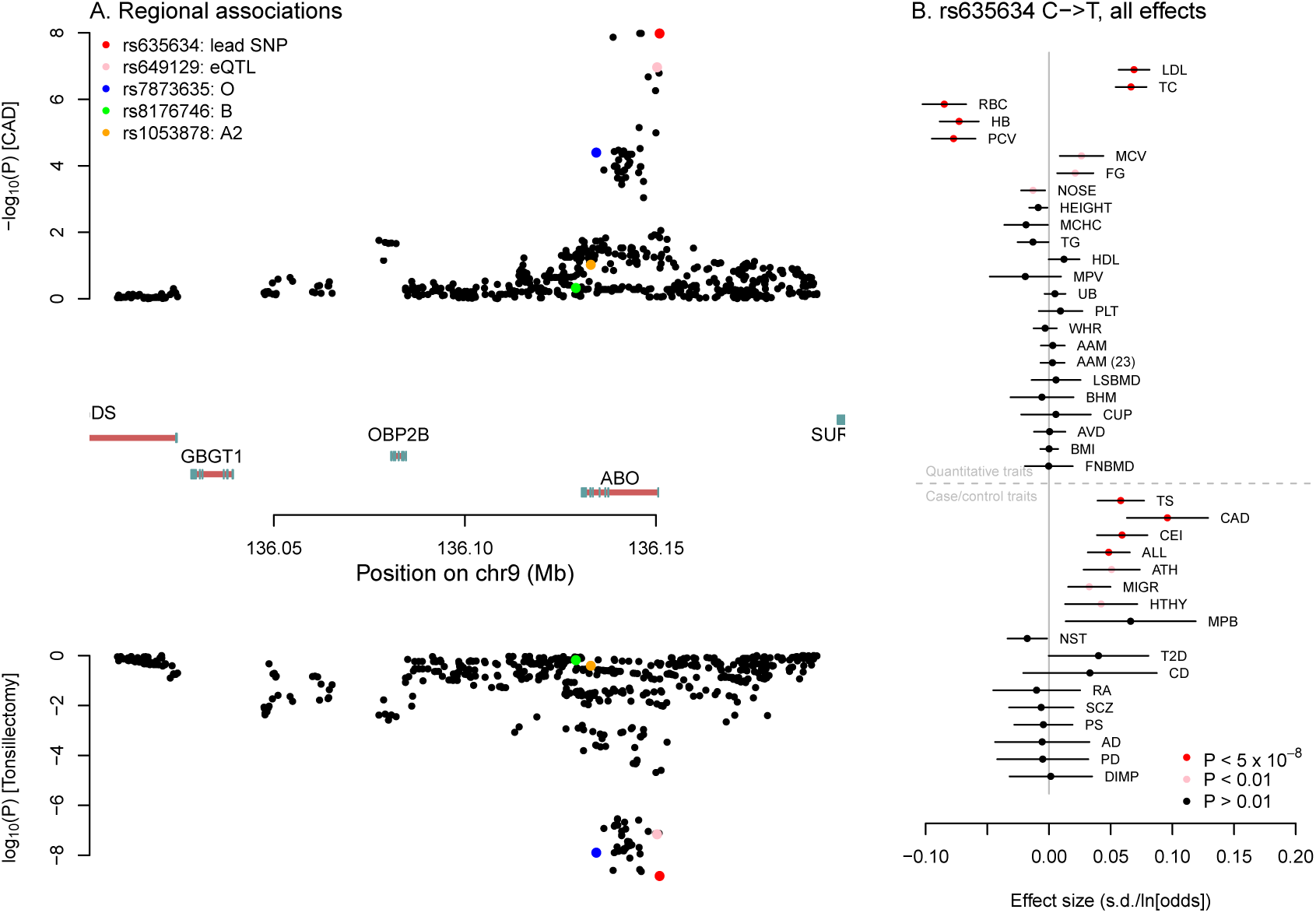
Multiple associations near the ABO gene. A. Association signals for coronary artery disease and tonsillectomy. In the top panel, we show the P-values for association with CAD for variants in the window around the ABO gene. In the bottom panel are the P-values for association with tonsillectomy. In both panels, SNPs that tag functionally-important alleles at ABO are in color. In the middle are the gene models in the region–exons are denoted by blue boxes, and introns with red lines. Note that the ABO gene is transcribed on the negative strand. **B. Association effect sizes for rs635634 on all tested traits.** Shown are the effect size estimates for rs635634 for all traits. The lines represent 95% confidence intervals. Traits are grouped according to whether they are quantitative traits (in which case the x-axis is in units of standard deviations) or case/control traits (in which case the x-axis is in units of log-odds).

Among the top-ranked regions are also a non-synonymous variant in the zinc transporter SLC39A8 (rs13107325; Supplementary Figure 9) that is associated with schizophrenia (log-odds ratio of the non-reference allele = 0.15, 95% CI = [0.11, 0.19], *P* = 2 *×* 10^−12^), Parkinson’s disease (log-odds ratio = - 0.15, 95% CI = [-0.21, -0.10], P = 1:6 × 10^−7^), and height (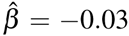 s.d., 95% CI = [-0.04, -0.02], *P* = 3.8 × 10^−7^), among others; a non-synonymous variant in the glucokinase regulator GCKR (rs1260326; Supplementary Figure 10) that is associated with fasting glucose 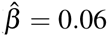 and height 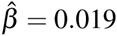, among others; and a region near the APOE gene (which we presume to be driven by the APOE4 allele; Supplementary Figure 11) that is associated with nearsightedness (log-odds ratio = -0.04, 95% CI = [-0.06, -0.02], *P* = 1.8 *×* 10^−5^), waist-hip ratio 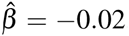, and several lipid traits apart from its well-known association with Alzheimer’s disease. It has previously been observed that association signals for different phenotypes tend to cluster spatially in the genome [Jeck et al., 2012]; these results suggest that in some cases clustered associations are driven by single variants. We note anecdotally that the variants that influence a large number of phenotypes seem to often be non-synonymous, rather than regulatory, changes, which contrasts with the pattern seen in association studies overall (e.g. Pickrell [2014]).

### 2.4 Identifying pairs of phenotypes with correlated effect sizes

In our scan for variants that influence pairs of phenotypes, we did not assume any relationship between the effect sizes of the variant on the two phenotypes. However, if two traits are influenced by shared underlying molecular mechanisms, we might expect the effect of a variant on the two phenotypes to be correlated. To test this, we returned to the set of variants identified by analysis of each phenotype individually (the numbers of these variants for each trait are in Table 1). For each set, we calculated the rank correlation between the effect sizes of the variants on the index trait (the one in which the variants were identified) and all of the other traits.

The results of this analysis are presented in Figure 4. Apart from closely related traits (e.g. the two measurements of bone density), we see a number of traits that are correlated at a genetic level. We focus on two of these. First, variants that delay age of menarche in women tend, on average, to decrease BMI (*ρ* = *−*0.53, *P* = 1.2 *×* 10^−6^, reduce risk of male pattern baldness (*ρ* = *−*0.45, *P* = 5.9 *×* 10^−5^), and increase height (*ρ* = 0.52, *P* = 2.2 *×* 10^−6^; Figure 4). These patterns hold both for the GWAS on age at menarche performed by Perry et al. [2014] and that performed by 23andMe (Figure 4). Most of these variants also delay age at voice drop in men (Figure 2), so we interpret these variants as ones that influence pubertal timing in general. The negative correlation between a variant’s effect on age at menarche and BMI has previously been observed [Bulik-Sullivan et al., 2015; Elks et al., 2010; Perry et al., 2014], as has the positive correlation between a variant’s effect on age at menarche and height [Bulik-Sullivan et al., 2015; Perry et al., 2014]. The negative correlation between a variant’s effect on age at menarche (or more likely, puberty in general) and male pattern baldness has not been previously noted, but is consistent with the known role for increased androgen signaling in causing hair loss [Hamilton, 1951; Li et al., 2012; Richards et al., 2008].

**Figure 4.**
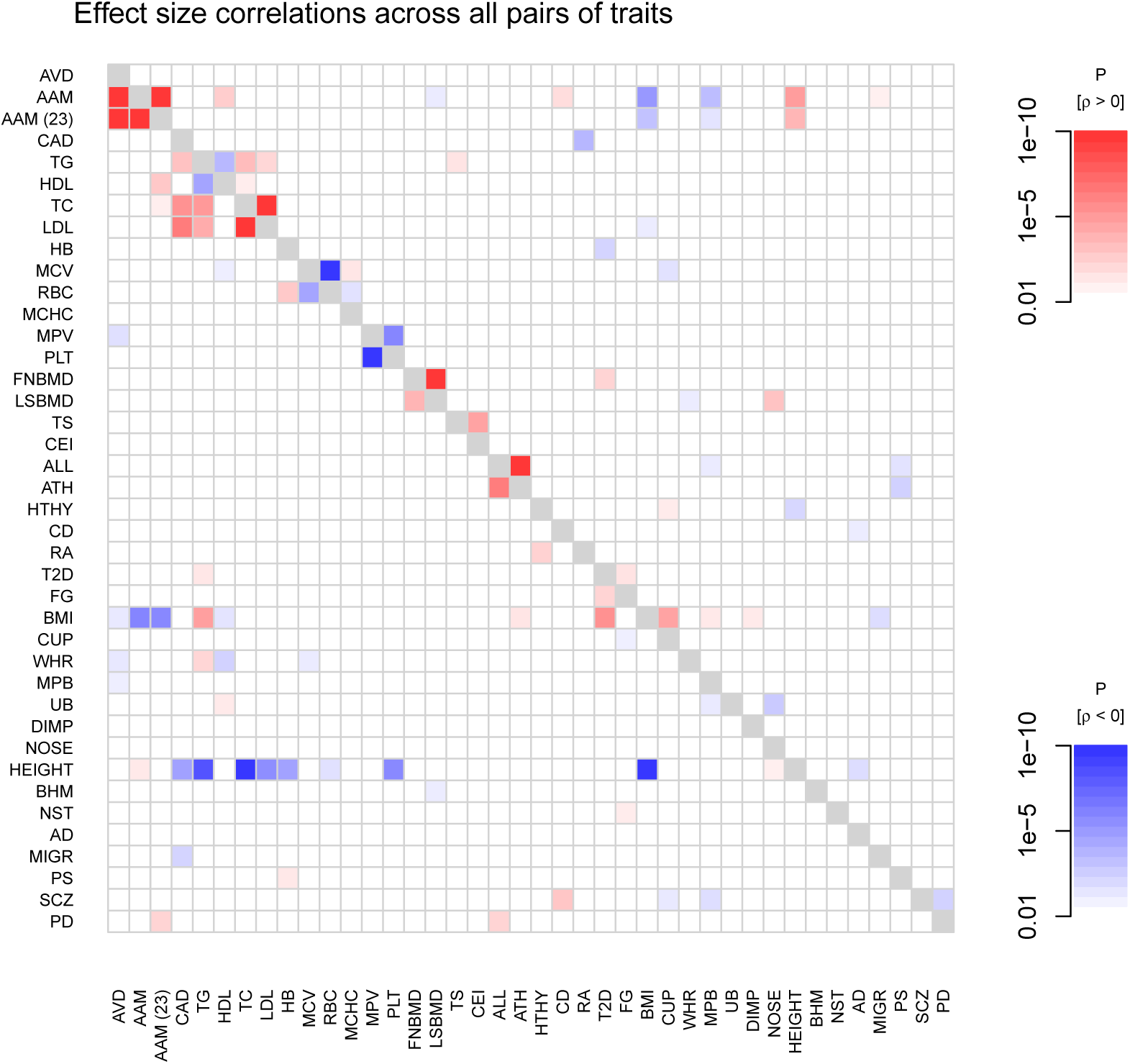
Heatmap showing patterns of correlated effect sizes of variants across pairs of traits. For each pair of traits [*i, j*], we extracted the set of variants that influence trait *i* and their effect sizes on both *i* and We then calculated Spearman’s rank correlation between the effect sizes on *i* and the effect sizes on *j*, and tested whether this correlation was significantly different from zero. Shown in color are all pairs where this test had a P-value less than 0.01. Darker colors correspond to smaller P-values, and the color corresponds to the direction of the correlation (in red are positive correlations and in blue are negative correlations). The phenotypes are in the same order as in Figure 2.

Second, we find that genetic variants that increase height tend to decrease triglycerides (*ρ* = *−*0.24, *P* = 4.2 *×* 10^−9^, LDL cholesterol (*ρ* = *−*0.2, *P* = 1.1 *×* 10^−6^), and risk of heart disease (*ρ* = *−*0.19, *P* = 6.4 *×* 10^−6^). These results are consistent with estimates of the genetic correlations between these traits [Bulik-Sullivan et al., 2015; Nelson et al., 2015] and the epidemiological observation that taller individuals have lower risk of heart disease [Gertler et al., 1951; Hebert et al., 1993; Paajanen et al., 2010]. Though the biological mechanism underlying this correlation is unclear (see discussion in Hebert et al. [1993] and Nelson et al. [2015]), the genetic data provides evidence against the existence of an unmeasured environmental confounding factor in the epidemiological studies.

### 2.5 Inferring causal relationships between traits

Finally, we were interested in identifying pairs of traits may be related in a causal manner. Since we are using observational data (rather than, for example, a randomized controlled trial), we view strong statements about causality as impossible. However, a realistic goal might be to identify aspects of the data that are more consistent with a causal model versus a non-causal model.

As a motivating example, we considered the correlation between levels of LDL cholesterol and risk coronary artery disease, now widely accepted as a causal relationship [Scandinavian Simvastatin Survival Study Group, 1994]. We noticed that variants ascertained as having an effect on LDL cholesterol levels have correlated effects on risk of coronary artery disease (Figure 4, Figure 5C), while variants ascertained as having an effect on CAD risk do not in general have correlated effects on LDL levels (Figure 5D). This is consistent with the hypothesis that LDL cholesterol is one of many causal factors that influence CAD risk. An alternative interpretation is that LDL cholesterol is highly genetically correlated to an unobserved trait that causally influences risk of CAD.

**Figure 5.**
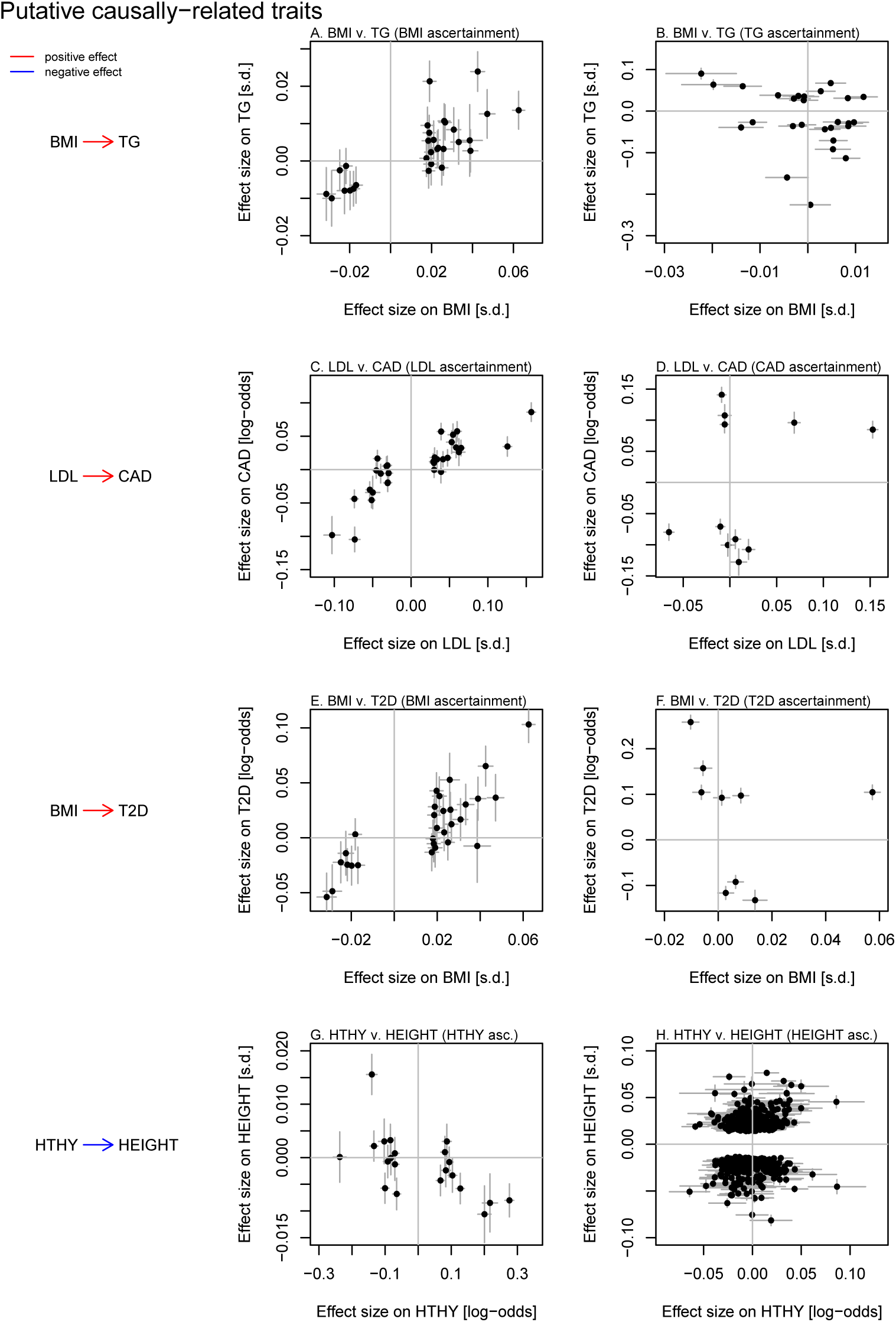
Putative causal relationships between pairs of traits. For each pair of traits identified as candidates to be related in a causal manner (see Methods), we show the effect sizes of genetic variants on the two traits (at genetic variants successfully genotyped or imputed in both studies). Lines represent one standard error. **A. and B. BMI and triglycerides.** The effect sizes of genetic variants on BMI and triglyceride levels for variants identified in the GWAS for BMI (**A.**) or triglycerides (**B.**). **C. and D. LDL and CAD.** The effect sizes of genetic variants on LDL levels and CAD for variants identified in the GWAS for LDL (**C.**) or CAD (**D.**). **E. and F. BMI and type 2 diabetes.** The effect sizes of genetic variants on BMI and T2D for variants identified in the GWAS for BMI (**E.**) or T2D (**F.**). **G. and H. Hypothyroidism and height.** The effect sizes of genetic variants on HTHY and height for variants identified in the GWAS for HTHY (**G.**) or height (**H.**).

We developed a method to detect pairs of traits that show this asymmetry in the effect sizes of associated variants, which we interpret as more consistent with a causal relationship between the traits than a non-causal one (Methods). At a threshold of a relative likelihood of 100 in favor of a causal versus a non-causal model, we identified five pairs of putative causally-related traits. (At a less stringent threshold of a relative likelihood of 20 in favor of a causal model, we identified 10 additional pairs of traits, see Supplementary Figure 10). Four of these are shown in Figure 4. First, genetic variants that influence BMI have correlated effects on triglyceride levels, while the reverse is not true; this suggests increased BMI is a cause for increased triglyceride levels (Figure 4). Randomized controlled trials of weight loss are also consistent with this causal link [Look AHEAD Research Group et al., 2007; Shai et al., 2008], as are Mendelian randomization studies [Freathy et al., 2008; Würtz et al., 2014]. Second, we confirm the evidence in favor of a causal role for increased LDL cholesterol in coronary artery disease (Figure 4), and in favor of a causal role for increased BMI in type 2 diabetes risk (Figure 4). Finally, we suggest that increased risk of hypothyroidism causes decreased height (Figure 4). While it is known that severe hypothyroidism in childhood leads to decreased adult height (e.g. Rivkees et al. [1988]), these data indicate that hypothyroidism susceptibility may also influence height in the general population. A fifth potentially causal relationship (between risk of coronary artery disease and rheumatoid arthritis) could not be confirmed in a larger study and so is not displayed (see Supplementary Information, Supplementary Figure 13).

## 3 Discussion

We have performed a scan for genetic variants that influence multiple phenotypes, and have identified several hundred loci that influence multiple traits. This style of scan complements methods to quantify the “genetic correlation” between two traits [Bulik-Sullivan et al., 2015; Lee et al., 2012; Loh et al., 2015; Visscher et al., 2014] that are not generally concerned with identifying individual variants that influence both traits. We were interested in using the individual variants identified to identify biological relationships between traits, including potential relationships when one trait is causally upstream of the other. Other potential mechanisms that could lead to an association between a genetic variant and two phenotypes include transgenerational effects of a variant on a parental phenotype and a separate phenotype in the offspring (e.g. Ueland et al. [2001]) or assortative mating that involves more than a single trait [Gianola, 1982].

**Genetic overlaps between traits.** One clear observation from these data is that genetic variants that influence puberty (age at menarche and age at voice drop) often have correlated effects on BMI, height, and male pattern baldness (Figure 4). In our scan for causal relationships between traits, we found modest evidence of a causal role of age at menarche in influencing adult height, and for a causal role of BMI in the development of male pattern baldness (Supplementary Figure 12). The non-causal alternative (also consistent with the data) is that all of these traits are influenced by some of the same underlying biological pathways, and perhaps the most likely candidate is hormonal signaling. This highlights the importance of considering evidence from multiple traits when interpreting the molecular consequences of a variant and designing experimental studies. While variants that influence height overall are enriched near genes expressed in cartilage [Wood et al., 2014] and variants that influence BMI are enriched near genes expressed broadly in the central nervous system [Locke et al., 2015], it seems a subset of these variants also influence age at menarche and male pattern baldness. For these variants, it may be worth considering functional follow-up in gonadal tissues or specific brain regions known to be important in hormonal signaling.

It is also striking to note how many genetic variants influence multiple traits (Figure 2) but without a consistent correlation in the effect sizes (Figure 4). For example, many of the autoimmune and immune-related traits appear to share many genetic causes in common, but the effect sizes of the variants on the different traits appear to be largely uncorrelated (see also Bulik-Sullivan et al. [2015]; Cotsapas et al. [2011]). Likewise, many variants appear to influence lipid traits, red blood cell traits and immune traits, but without consistent directions of effect. A trivial explanation of this observation is that we are underpowered to detect correlations in the effect sizes because we are using only a small set of the SNPs with the strongest associations. However, the genetic correlations between many of these traits (calculated using all SNPs) are not significantly different from zero [Bulik-Sullivan et al., 2015]. Another possibility is that a given genetic variant often influences the function of multiple cell types through separate molecular pathways, or that the effects of a variant on two related phenotypes vary according to an individual’s environmental exposures.

**Causal relationships between traits.** From the point of view of epidemiology, the ability to scan through many pairs of traits to find those that are potentially causally related seems appealing, and some previous analyses have had similar goals [Evans et al., 2013]. Our approach makes the key assumption that, if two traits are related in a causal manner, then the “causal” trait is one of many factors that influence the “caused” trait. This induces an asymmetry in the effects of genetic variants on the two traits that can be detected (Figure 5). We also assume that we have identified a modest number of variants that influence both traits. This naturally means we are limited to considering heritable traits that have been studied with in cohorts with moderate sample sizes (on the order of tens to hundreds of thousands of individuals). It seems likely that the main limiting factor to scaling this approach (should it be generally useful) will be phenotyping rather than genotyping.

## 4 Methods

We obtained summary statistics from genome-wide association studies described in Table 1. The complete description of these studies is in the Supplementary Materials.

### 4.1 Counting independent numbers of associated variants

For each genome-wide association study, we ran fgwas v.0.3.6 [Pickrell, 2014] with the default settings, except that rather than splitting the genome into blocks with equal numbers of SNPs (as in Pickrell [2014]), we split the genome into approximately independent blocks based on patterns of linkage disequilibrium in the European populations in Phase 1 of the 1000 Genomes Project [Abecasis et al., 2010]. These blocks are available at https://bitbucket.org/nygcresearch/ldetect-data. We used fgwas to estimate the prior probability that any block contains an association. The output of this model is, for each region of the genome, the posterior probability that it contains a variant that influences the trait. We used a threshold of a posterior probability of association of 0.9, as in Pickrell [2014], which can be roughly interpreted as a false discovery rate of 10%. For analyses that use variants identified in these individual GWAS, we extracted the single SNP from each region with the largest posterior probability of being the causal SNP in this model.

### 4.2 Approximating the correlations in the effect sizes under the null model

For genome-wide association studies of correlated traits performed on overlapping individuals, we expect the observed association statistics of a given variant to both traits to be correlated, even under the null model that the variant influences neither trait. To approximate this expected correlation, for both traits we extracted all genomic regions with a posterior probability of containing an association less than 0.2 (using the method described above). We then extracted all SNPs from these regions, and calculated the correlation in the Z-scores between the two traits (using all SNPs remaining in both studies). This correlation is a function of the number of overlapping samples and the correlation in the phenotypes. Specifically, if 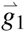 is a vector of (mean-centered) genotypes at a variant in study 1, 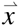 is a vector of (standard normally distributed) phenotypes in study 1, 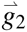 is a set of is a vector of genotypes at a variant in study 2, and 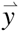 is a vector of phenotypes in study 2, then:

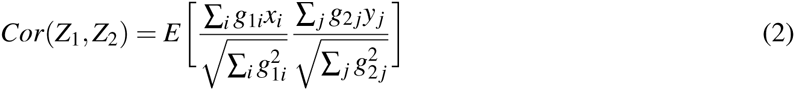

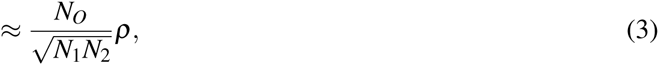

where *N*_*O*_ is the number of overlapping individuals in the two studies, *N*_1_ is the number of individuals in study 1, *N*_2_ is the number of individuals in study 2, and *ρ* is the correlation between the phenotypes. We used this correlation as a correction factor in all pairwise GWAS.

### 4.3 Hierarchical model

In this section we describe the hierarchical model used for the main scan for overlapping association signals in two GWAS. Our goal is to write down a model that allows us to estimate the probability that a genomic locus contains a variant that influences two traits. Our approach is to split the genome into non-overlapping regions (we used the same approximately independent blocks as above); each region then falls into one of five categories, following Giambartolomei et al. [2014]:

0. There are no SNPs in the region that influence either trait (denoted *RM*_0_, for regional model 0),
1. There is one causal SNP in the region that influences the first trait (*RM*_1_),
2. There is one causal SNP in the region that influences the second trait (*RM*_2_),
3. There is one causal SNP in the region that influences both traits (*RM*_3_),
4. There are two causal SNPs in the region, one of which influences the first trait and one of which influences the second (*RM*_4_).

We will estimate the proportion of genomic regions in each of these categories with an empirical Bayes approach. In what follows, we start by writing down the model for the simplest case where two phenotypes have been studied in separate cohorts, and then introduce modifications for the more complex situations that arise in real data. Software implementing the model is available at https://github.com/joepickrell/gwas-pw.

In the simplest case, consider two separate genome-wide association studies performed on two traits. In this case, the model is a hierarchical version of that in Giambartolomei et al. [2014], which we re-iterate here for completeness. Let there be *N*_1_ individuals in GWAS of the first phenotype and *N*_2_ individuals in the GWAS of the second phenotype. We start by considering a single SNP. Let 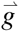 be the vector of genotypes at the SNP in the first study, 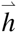 be the vector of genotypes at the SNP in the second study, 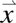 be the vector of phenotype measurements for the first phenotype (assumed to be distributed as a standard normal), and 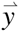 be the vector of phenotype measurements for the second phenotype (also assumed to be distributed as a standard normal). We first need a measure of the evidence that the SNP influences each of the traits.

#### 4.3.1 Bayes factor calculations

We use a simple linear regression model to relate the phenotypes and the genotypes:

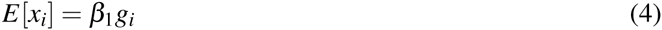

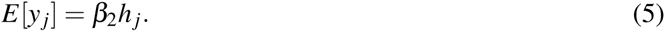

There are four potential models to consider for this SNP:

0. *M*_0_: the SNP is associated with neither trait
1. *M*_1_: the SNP is associated with the first trait (but not the second)
2. *M*_2_: the SNP is associated with the second trait (but not the first)
3. *M*_3_: the SNP is associated with both traits.

Model *M*_0_ corresponds to the case where *β*_1_ = 0 and *β*_2_ = 0, model *M*_1_ corresponds to the case where *β*_1_ is free to vary while *β*_2_ = 0, and so on. We can thus define three Bayes factors corresponding to the evidence in favor of the three alternative models:

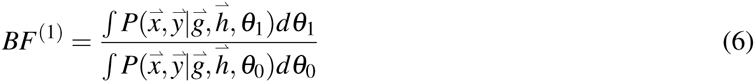

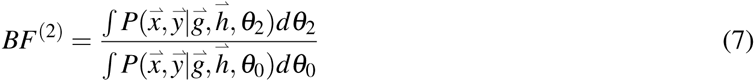

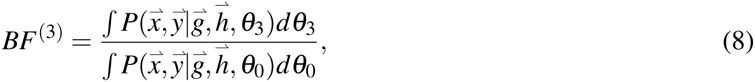

where *θ*_*j*_ represents the parameters of model *j*. To compute these Bayes factors, we use the approximate Bayes factors from Wakefield [2008]. If we let 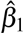 be the maximum likelihood estimate of *β*_1_ and 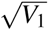 be the standard error in that estimate, then 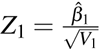. If the prior on the true effect size is *β*_1_ *∼ N*(0,*W*_1_) we can write down the Wakefield approximate Bayes factor measuring the evidence that the SNP is associated with the first phenotype:

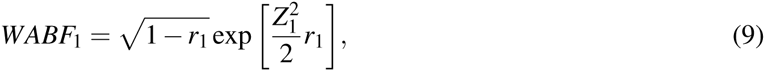

where 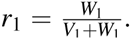 *WABF*_2_ is defined analogously. In all applications, we averaged over Bayes factors computed with *W* = 0.01,*W* = 0.1, and *W* = 0.5. To now connect these approximate Bayes factors to the three alternative models for the SNP [Giambartolomei et al., 2014]:

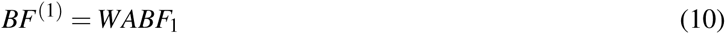

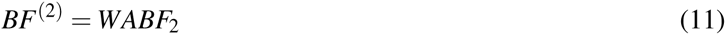

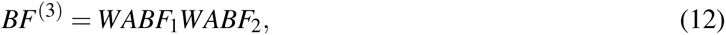

where *BF*^(3)^ is a consequence of the fact that the two cohorts are independent. This latter Bayes factor is equivalent to that derived under the “maximum heterogeneity” model in Wen and Stephens [2014].

We note that in the Wakefield approximate Bayes factor the effect size of a SNP enters only through the Z-score. As a consequence, if we consider the “reverse” regression model where we swap the genotypes and phenotypes:

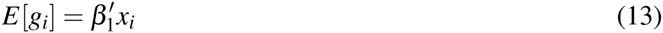

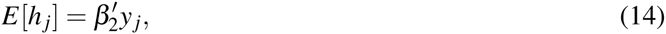

then the Bayes factors from this model are identical to the previous model (as long as the ratios *r*_1_ and *r*_2_ remain constant). In fact it is this latter “reverse” regression that we will use going forward, though it is simpler to interpret parameters like the prior on the effect size in the traditional parameterization.

#### 4.3.2 Regional Bayes factor

We now consider a Bayes factor measuring the support for an association in a given genomic region *r*. To do this, now consider the matrix **G**_*r*_ of genotypes in the region in the first study (with *N*_1_ rows of individuals and *K* columns of SNPs) and the matrix **H**_*r*_ of genotypes in the region in the second study (with *N*_2_ rows and *K* columns). The vectors of phenotypes remain 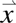 and 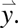 We now want to write down Bayes factors measuring the evidence in favor of the four alternative models discussed at the beginning of Section 4 relative to the null model of no associations in the region. For regional model 1 (there is a single SNP casually associated with the first phenotype and none with the second phenotype):

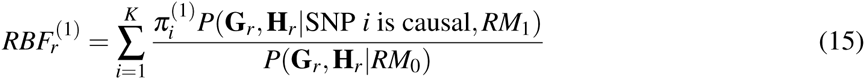

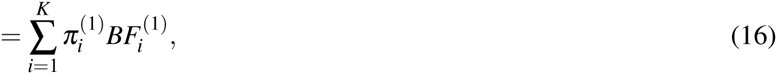

where 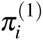 is the prior probability that SNP *i* is the causal one under model 1. Note that the probabilities of all genotypes at the non-causal sites cancel out because they are identical once we have conditioned on the genotype at a causal site [Maller et al., 2012].

Analogously,

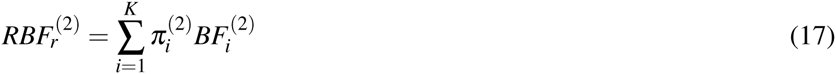

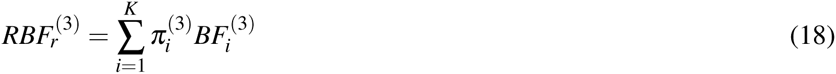

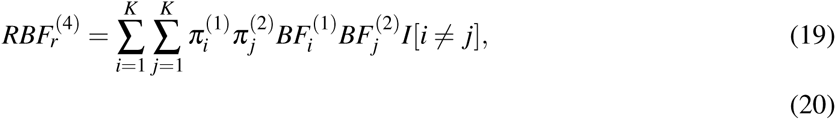

where *I*[*i* ≠ *j*] is an indicator that evaluates to 1 if *i* and *j* are different and 0 otherwise, 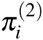 is the prior probability that SNP *i* is the causal SNP under model 2, and 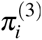 is the prior probability that SNP *i* is the causal SNP under model 3. For model 4 (where there are two causal SNPs, one of which only influences the first phenotype and one of which only influences the second phenotype), we assume that the prior probabilities that SNP *i* influences the first or second phenotype are identical to those under model 1 and 2, respectively. In all applications we set 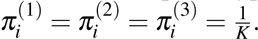.

#### 4.3.3 Likelihood

We now turn to the model for the whole genome. We denote the full matrix of genotypes in the first study as **G** and the full matrix of genotypes in the second study as **H**. We split the genome into *M* approximately independent blocks. Under the assumption that all blocks are independent, the probability of all the genotypes is:

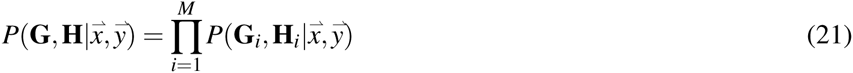

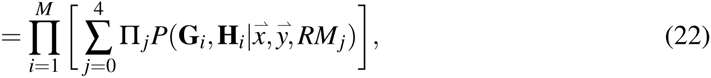

where Π _*j*_ is the prior probability of regional model *j* and *RM*_*j*_ is regional model *j*. These are the probabilities we would like to learn. We can do so by maximizing the log-likelihood:

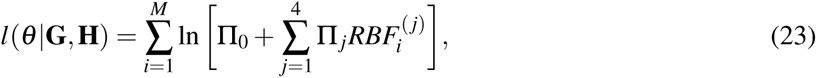

where *θ* is the set of parameters in the model (the prior probabilities and all of the parameters that go into the construction of the Bayes factors). We maximized this likelihood with the approach described in Section 4.3.5.

#### 4.3.4 Bayes factors for overlapping cohorts

The above model makes the key assumption that the two phenotypes in question have been measured on two separate sets of individuals. In practice, the cohorts we will use are often overlapping or partially overlapping. This causes two problems for the model. First, if the two phenotypes are correlated, we may overestimate the evidence in favor of regional model 3 (where a single variant influences both phenotypes). Second, the patterns of linkage disequilibrium in the population can no longer be ignored when considering the evidence in favor of regional model 4, and we may overestimate the evidence in favor of this model. In practice, we were most concerned with the first of these; see the Supplementary Material for discussion of the second.

The degree to which we may overestimate the evidence in favor of regional model 3 depends on the number of overlapping samples in the two studies and the correlation in the phenotypes. We consider the case where there is a single cohort of individuals. Let the vector of genotypes at the SNP be 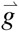 and let the two vectors of phenotypes be 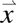 and 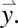 Let the phenotypes be bivariate-normally distributed with mean zero, variance one, and correlation coefficient *C*.

As before, we first want to calculate the Bayes factors measuring the evidence in favor of the three alternative models from Section 4.3.1. We use a multivariate linear regression model:

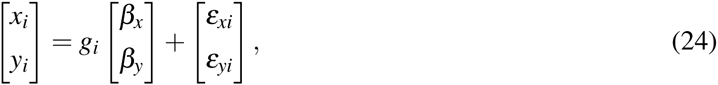

where *β*_*x*_ is the effect of the SNP on phenotype *x*, *β*_*y*_ is the effect of the SNP on phenotype *y*, and *ε*_*xi*_ and *ε*_*yi*_ are error terms that are multivariate normally distributed with mean zero and covariance matrix Σ (though in all that follows we assume the effects of any SNP are small, so this residual covariance matrix is approximated by the covariance matrix of the phenotypes).

To compute the Bayes factors, we use a multivariate extension of the approximate Bayes factor from Wakefield [2008]. Instead of working directly with the phenotype and genotype vectors, we instead consider 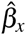 and 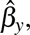 the estimated effect sizes from each individual regression. We also use *V*_*x*_ and *V*_*y*_, the respective variances in the estimates of each regression coefficient, and 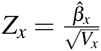 and 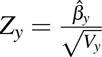. We let:

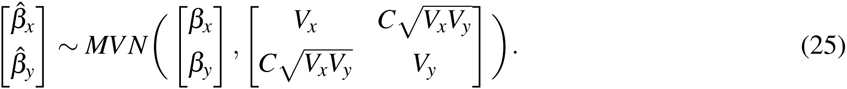

We now place a multivariate normal prior on *β*_*x*_ and *β*_*y*_. The form we choose is:

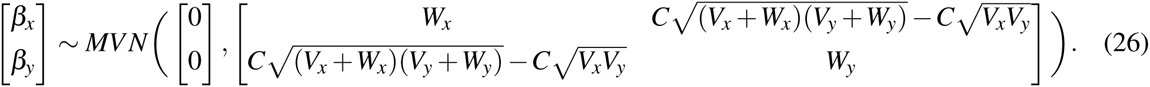

This prior has the somewhat odd property that it depends to a small extent on the variances of the effect size estimates (i.e. on the minor allele frequency of the SNP in question), such that rarer SNPs, or those with a large amount of missing data, have larger prior covariances. The benefit of this prior is that now the posterior predictive distribution of the estimated effect sizes has a simple form:

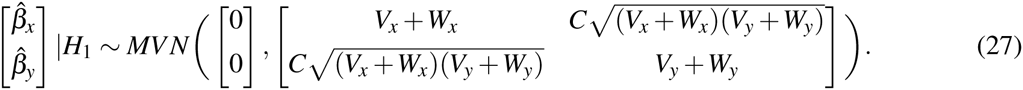

The other models are similar. With these assumptions, we can analytically compute the Bayes factors:

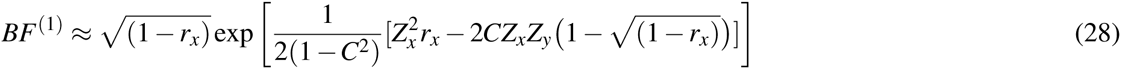

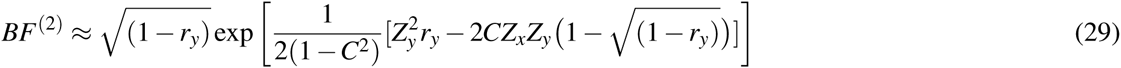

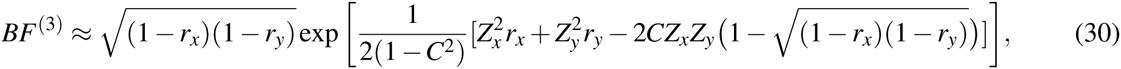

where 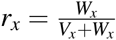 and 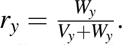 Note that if the two phenotypes are uncorrelated all three Bayes factors are identical to those in Section 4.3.1.

For all of the pairwise GWAS, we used these Bayes factors instead of those in Section 4.3.1. We used *C* estimated from the summary statistics, as described in Section 4.2. Note that when the cohorts are only partially overlapping, the *C* we calculate is a function of the amount of overlap between the cohorts as well as the correlation in the phenotypes. In principle some knowledge of the true correlation between two phenotypes could be obtained from external data and incorporated into the prior here, but we have chosen not to do this, and so as the overlap in the cohorts goes to zero, these Bayes factors tend to the prior assumption that the two phenotypes are uncorrelated (i.e. the maximum heterogeneity Bayes factor from Wen and Stephens [2014]). This has no justification from a modeling perspective, and so may be suboptimal in situations where good external information is available.

#### 4.3.5 Fitting the model

The natural approach to fitting this model would be to maximize the log-likelihood in Equation 23. However, in a small subset of cases (generally in pairs of GWAS with small numbers of associated variants), we found that this maximization was numerically unstable. To fix this, we placed a weak logistic normal prior on the Π parameters. Specifically, we define hyperparameters:

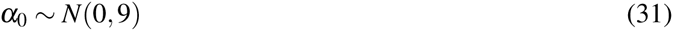

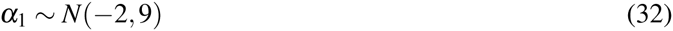

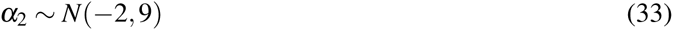

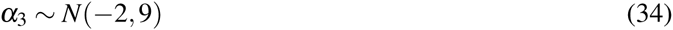

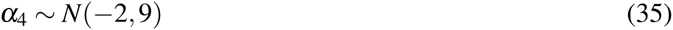

and then define:

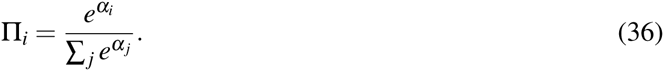

Instead of maximizing the likelihood, we maximized the function:

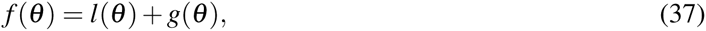

where *θ* is the set of five *α* parameters, *l*(*θ*) is the log-likelihood from Equation 23 and *g*(*θ*) is the log of the prior density described above. We maximized this function using the Nelder-Mead algorithm implemented in the GNU Scientific Library. The estimates of the parameters are maximum *a posteriori* estimates rather than maximum likelihood estimates. In practice, this serves to prevent estimates of the Π parameters from going all the way to zero.

### 4.4 More robust “Mendelian randomization”

The observation that a genetic variant influences two traits can be interpreted in the “Mendelian randomization” framework as evidence that one trait causally influences the other. However, using this framework requires the strong assumption that the variant does not influence the two traits via independent mechanisms.

Our goal was to develop a robust method for measuring the evidence in favor of a causal relationship between two traits using data from many genetic associations, while recognizing that strong conclusions are likely impossible in this setting. Our aim was not to estimate the magnitude of a causal effect (should one exist), but rather to simply determine if such an effect exists.

Our motivating example comes from LDL cholesterol and heart disease risk–if we identify variants that influence LDL levels, these variants have correlated effects on heart disease risk (Figure 5). However, if we identify genetic variants that influence heart disease, these variants do not have correlated effects on LDL levels (Figure 5). The intuition is as follows: if a trait *X* causally influences trait *Y*, then to a first approximation every genetic variant that influences trait *X* should also influence trait *Y*, and the effect sizes of these variants on the two traits should be correlated. The reverse, however, is not true: genetic variants that influence trait *Y* do not necessarily influence trait *X*, since *Y* can be influenced by mechanisms independent of *X*. (Assume that *X* is one of a large number of factors that causally influence *Y*, such that most of the variants that influence *Y* do not act through *X*).

To scan through all pairs of traits, we aimed to formulate this intuition in a manner that allows for automation. Related work has been done on Mendelian randomization with multiple genetic variants [Davey Smith and Hemani, 2014; Do et al., 2013; Evans et al., 2013] and “reciprocal” Mendelian randomization [Timpson et al., 2011]. We assume we have identified a set of *N*_*X*_ genetic variants that influence *X* (without using information about *Y*). Assume we have also identified a set of *N*_*Y*_ genetic variants that influence *Y* (without using information about *X*). Let 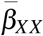 be the vector of effect sizes on trait *X* for the set of variants ascertained through the association study of *X*, and 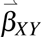 be the vector of the effect sizes of these variants on trait *Y*. Define 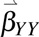 and 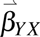 analogously. Now let 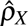 be the rank correlation between 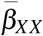 and 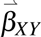, and let 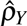 be the rank correlation between 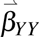 and 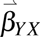. Using Fisher’s Z-transformation, we can approximate the sampling distributions of 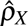 and 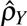. If we let 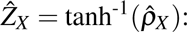

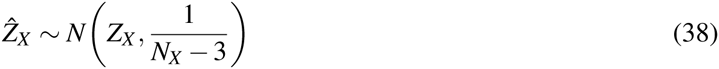

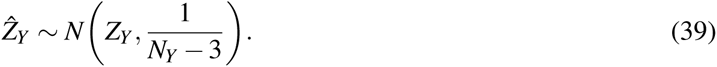

We can thus define an approximate likelihood for the two correlation coefficients:

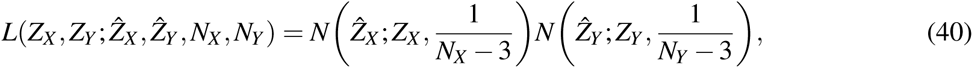

where *N*(*x*; *μ*, *σ*^2^) is the density of a normal distribution with mean *μ* and variance *σ*^2^ evaluated at *x*.

Now we define the models we would like to compare:

1. M1: if trait *X* causes *Y*, then we estimate *Z*_*X*_ and set *Z*_*Y*_ = 0.
2. M2: If trait *Y* causes *X*, then we estimate *Z*_*Y*_ and set *Z*_*X*_ = 0.
3. M3: If there are no relationships between the traits, then *Z*_*X*_ = *Z*_*Y*_ = 0.
4. M4: If the correlation does not depend on how the variants were ascertained, *Z*_*X*_ = *Z*_*Y*_.

The first two models on this list we think of as causal models, by the line of reasoning outlined at the beginning of this section. The third model we obviously interpret as a non-causal model. The fourth we also interpret as non-causal, though this perhaps merits some discussion (see below). We fit each model by maximizing the corresponding approximate likelihood.

To compare the models, we calculate the Akaike information criterion (AIC) for each, where the numbers of parameters is 1, 1, 0, and 1, respectively, for the four models above. We then choose the smallest AIC from the two causal models (*AIC*_*causal*_) and the smallest AIC from the two non-causal models (*AIC*_*noncausal*_). We then calculate the relative likelihood of these two models:

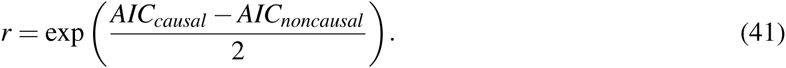

This is the relative likelihood of the best non-causal model compared to the best causal model. In Figure 5, we show the four pairs of traits where this ratio is less than 0.01, and in Supplementary Figure 12, we show 10 additional pairs of traits where this ratio is less than 0.05. As we can see visually, this model successfully identifies patterns that look similar to our motivating example of LDL and heart disease.

A key caveat in interpretation of this method is that we may not have measured the truly causal phenotype, but rather some proxy for it. For example, if it is not BMI *per se* that causally influences risk of type 2 diabetes, but rather some other measure of adiposity that is highly correlated to BMI (and shares the same underlying genetic basis), then we have no way of detecting this. We suspect that more detailed phenotyping will identify “clusters” of highly correlated traits that will be difficult to disentangle.

**Implications of looking explicitly for asymmetry.** We have set up this model in the context where a causal trait is one of many factors that influences a downstream trait. This induces the asymmetry we try to detect. However, it is possible that the “causal” trait is the *major* factor that influences the “caused” trait. For example, consider type 2 diabetes and fasting glucose levels. Clearly any factor that increases fasting glucose increases risk of type 2 diabetes, just by virtue of the definition of the disease. This type of causal relationship will be missed by this approach. On the other hand, consider the two measures of bone mineral density. Any factor that increases one will also almost certainly increase the other, because the two phenotypes are closely related at a molecular level. We would not consider this a causal relationship between the traits, but rather that the two traits are measurements of a single underlying variable (namely overall bone density). We prefer to miss causal relationships of the first kind in order to avoid the interpretive difficulties of the second case.

## 5 Acknowledgements

This work was supported in part by the National Human Genome Research Institute of the National Institutes of Health (grant number R44HG006981 to 23andMe) and the National Institute of Mental Health (grant number R01MH106842 to JKP). We would like to thank the customers of 23andMe for making this work possible, the GWAS consortia that made summary statistics available to us, Luke Jostins for providing updated summary statistics from the Crohn’s disease GWAS, and Graham Coop and Matthew Stephens for helpful discussions.

